# Technology applications in bovine gait analysis: a scoping review

**DOI:** 10.1101/2022.03.18.484936

**Authors:** Amir Nejati, Anna Bradtmueller, Elise Shepley, Elsa Vasseur

## Abstract

Quantitative bovine gait analysis using technology has evolved significantly over the last two decades. However, subjective methods of gait assessment using visual locomotion scoring remain the primary on-farm and experimental approach. The objective of this review is to map research trends in quantitative bovine gait analysis and to explore the technologies that have been utilized to measure biomechanical parameters of gait. A scoping literature review was conducted according to PRISMA guidelines. A search algorithm based on PICO framework generated three components – bovine, gait, and technology – to address our objectives. Three online databases were searched for original work published from January 2000 to June 2020. A two-step screening process was then conducted, starting with the review of article titles and abstracts based on inclusion criteria. A remaining 125 articles then underwent a full-text assessment, resulting in 82 final articles. Thematic analysis of research aims resulted in four major themes among the studies: gait/claw biomechanics, lameness detection, intervention/comparison, and system development. Lameness detection (55 % of studies) was the most common reason for technology use. Studies in the field of bovine gait analysis used three main technologies: force and pressure platforms (FPP), vision-based systems (VB), and accelerometers. FPP were the first and most popular technologies to evaluate bovine gait and were used in 58.5 % of studies. They include force platforms, pressure mapping systems, and weight distribution platforms. The second most applied technology was VB (34.1 % of studies), which predominately consists of video analysis and image processing systems. Accelerometers, another technological method to measure gait characteristics, were used in 14.6 % of studies. A strong demand for automatic lameness detection influences the path of development for quantitative gait analysis technologies. Although progress has been made, more research is needed to achieve more accurate, practical, and user-friendly technologies.

## Introduction

Gait abnormalities are a major welfare concern that cause considerable economic losses in cattle farming (1, 2). Hoof and leg disorders are the most common causes of gait abnormalities – mainly referred to as “lameness” in literature (3, 4). Identifying cows with impaired locomotion, especially at early stages of development, is key to minimizing the welfare and economic consequences of gait abnormalities (5). The prevalence of lame cows is often underestimated by farmers who only rely on passive observation of cows for abnormal gait (e.g., visual assessment when moving cows for milking), which is not sufficient for identifying lame cows (6), particularly in the case of mild lameness and early lameness detection. Gait analysis can be used to facilitate and improve lameness detection in cows (7). Besides the role of gait analysis in identifying a lame cow, it can also be utilized in research to investigate cow’s welfare and ease of movement under different conditions (e.g., housing systems, Shepley and Vasseur (8); flooring type, Telezhenko and Bergsten (9)).

Generally, gait analysis is conducted through subjective (qualitative) and objective (quantitative) methods. Subjective methods, i.e., visual gait scoring, is the traditional and primary method of gait analysis as it is easy and inexpensive to implement both on-farm and in research studies. However, visual gait scoring is subjective in nature and, as such, variability in the observer’s training level and background experience as well as the wide range of different gait characteristics evaluated across different scoring systems may contribute to low intra- and inter-observer reliability (3, 10, 11). In addition, visual gait scoring is a time-consuming procedure, particularly in large herds, which may lead to producers conducting less frequent assessments of gait, resulting in later detection of lameness and, thus, hindering early intervention and treatment (3). Objective methods using gait analysis technologies have been developed by researchers over the years to address the weaknesses of the traditional methods of cow gait assessment. However, exploring and appraising studies that utilize gait analysis technologies and drawing comparisons between their outcomes is difficult. This is due not only to the variety of ways in which gait analysis technologies are applied in research and the outcome measures investigated, but also due to differences in hardware and settings within similar types of technology that complicate comparisons even when the same category of gait analysis technology is used. Moreover, although several efforts have been made to put forth systematic reviews on cow gait analysis technologies, these looked primarily at studies in which cow lameness is the central focus. There is a lack of systematic reviews that considers all aspects of technology-based cow gait and movement analysis, independent of cow lameness.

The objectives of this scoping review were 1) to map research trends of quantitative bovine gait analysis, 2) to explore the technologies that have been being utilized to measure biomechanics parameters of gait variables in bovine species, and 3) to spotlight the current gaps in the field of cow gait analysis.

## Methods

The protocol for this review was formatted as per the items for Systematic Reviews and Meta-Analyses extension for Scoping Reviews, i.e., PRISMA-ScR (12).

### Eligibility criteria

English language full text publications of primary research from any geographic location were included without restrictions on study design. Publications dated prior to 2000 were excluded due to substantive changes that occurred since that time in bovine gait practices. We included research conducted on live cow populations (in vivo studies) for either beef or dairy cattle with no age and production cycle limitation. Only studies that measured gait attributes objectively through technologies using biomechanical parameters derived from kinematic and kinetic analysis were included in the review.

### Information sources and search strategy

To identify references, the literature search was conducted in three electronic databases (Web of Science Core Collection, CAB Abstracts, and Scopus). A comprehensive search strategy was developed in order to identify relevant literature. Search terms were developed based on PICO framework in which three main components-population/problem (P), intervention/interest (I), context/comparison (Co)- were extracted from the original research question. The target population (first component) is bovine species in which we are looking for the intervention of technology (second component) in the context of gait analysis (third component). Afterwards, nine domains were included in the search. These domains were bovine, gait analysis, locomotion analysis, movement analysis, lameness detection, biomechanics, vision-based analysis, accelerometer, and measuring plates. The search algorithm applied the following combination of these nine domains: (bovine) AND [(gait analysis) OR (locomotion analysis) OR (movement analysis) OR (lameness detection)] AND [(biomechanics) OR (vision-based analysis) OR (accelerometer) OR (measuring plate)]. Then, in the selected databases, article titles, abstracts and keywords using the nine domains with several possible keywords for each were looked at. Table 1 shows the details of the search strings conducted in the Web of Science database as an example. The same search strategy was translated into the CAB Abstracts and Scopus databases. The final search was conducted on June 2, 2020. For this search, no limits were set on language, subject area, study design or date of publication to allow for the minimization of bias in identifying all relevant research for inclusion in the review.

**Table 1.** List of the different strings included in the search strategy and the number of retrieved references for each string. Application on the Web of Science database on June 2, 2020.

| # | String | Records found |
| --- | --- | --- |
| # 1 | TS=(cow\$ OR bovine OR cattle OR heifer\$ OR calf OR calves) | 595,051 |
| # 2 | TS=((movement NEAR/15 analys*) OR (movement NEAR/15 evaluat*) OR (movement NEAR/15 measur*) OR (movement NEAR/15 assess*)) | 42,272 |
| # 3 | TS=((locomot* NEAR/15 analy*) OR (locomot* NEAR/15 evaluat*) OR (locomot* NEAR/15 measur*) OR (locomot* NEAR/15 assess*)) | 13,507 |
| # 4 | TS=((gait NEAR/15 analy*) OR (gait NEAR/15 evaluat*) OR (gait NEAR/15 measur*) OR (gait NEAR/15 assess*)) | 9,519 |
| # 5 | TS=((lameness NEAR/15 detect*) OR (lameness NEAR/15 evaluat*) OR (lameness NEAR/15 measur*) OR (lameness NEAR/15 identif*)) | 1,252 |
| # 6 | TS=(kinesi* OR kinetic\$ OR kinemat* OR biomechanic\$) | 595,051 |
| # 7 | TS=(acceler* OR sensor\$) | 1,451,335 |
| # 8 | TS=((force NEAR/15 (plate\$ OR mat\$)) OR (sensitive NEAR/15 (plate\$ OR mat\$)) OR (pressure NEAR/15 (plate\$ OR mat\$)) OR GRF OR (ground NEAR/15 force)) | 14,549 |
| # 9 | TS=(3\$d OR 3\$dimension* OR three\$dimension* OR 2\$d OR 2\$dimension* OR two\$dimension* OR (video\$ NEAR/15 (process* OR analy* OR based)) OR (image\$ NEAR/15 (process* OR analy* OR based)) OR (vision NEAR/15 (based OR computer))) | 586,120 |
|  | #1 AND (#2 OR #3 OR #4 OR #5) AND (#6 OR #7 OR #8 OR #9) | 280 |

A supplementary search was performed using hand-searching to pick up any relevant references that were missed by the database searches. Subsequently, the reference lists of each of the documents found to meet the inclusion criteria were also screened to identify any additional documents of interest.

### Selection of sources of evidence

All references were imported into the Endnote X9 reference management software (13) and duplicates were removed. All literature was then uploaded to Covidence, an online systematic review management program (14). The review was then performed in two steps. The first consisted of a review of titles and abstracts through which papers unrelated to the research questions (n=349) were excluded. Articles related to the topic of bovine gait and movement analysis or lameness detections were all included in the next step of screening process. The second step consisted of a full-text review to make sure that the references all met the eligibility criteria. Further exclusion of the documents was performed during the data collection process (i.e., a document could be later excluded based on its full-text review). Fig 1 summarizes the selection process.

**Fig 1.**
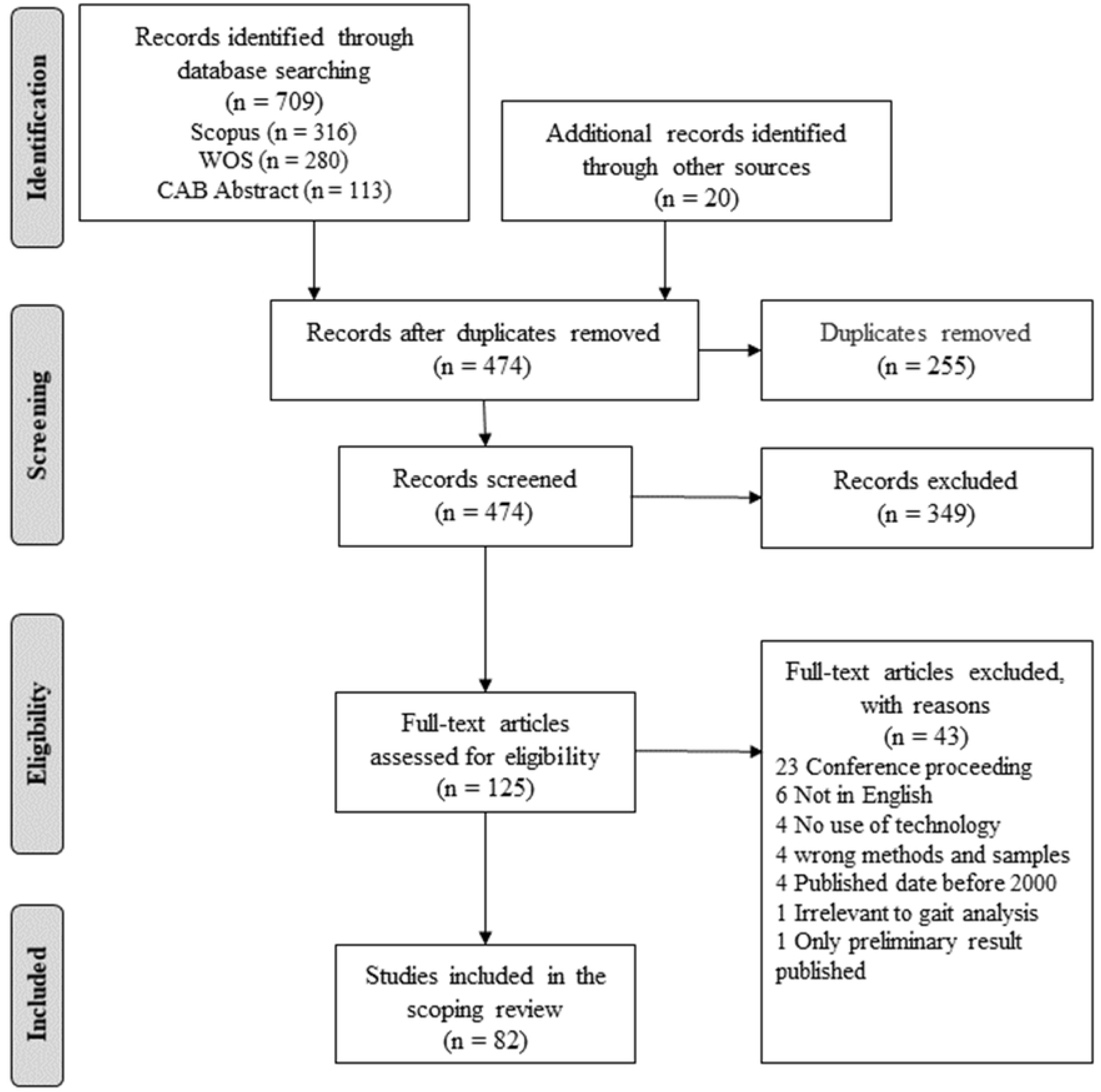
PRISMA flow diagram for scoping reviews showing literature search and selection of articles.

### Data charting process and data items

A data extraction sheet developed by the authors was used to chart literature under the following headings: author(s), title, year published, journal, research aims, research setting (laboratory- or farm-based), technologies used, hardware (device, specifications), types of animal preparation (e.g., marker or sensor attachment), target anatomical regions, test corridor characteristics (dimensions, flooring, number of steps), housing system characteristics (e.g., housing type, flooring type), sample size, breed, age, production cycle (dry or lactating), measured variables, main findings, study limitations, and future directions. All screening and data extraction steps were performed by a single reviewer. To minimize the likelihood of human error, all the uncertainties during the review process as well as the initial development of the scoping review protocol were discussed with the review team on a weekly basis.

### Synthesis of results

A narrative synthesis was used to collate, summarize, and present the findings of the current review. Various tools and approaches such as thematic analysis, textual descriptions, tabulation, and graphs were used to explore similarities, differences, and relationships between different studies. Literature data were analyzed using a thematic analysis approach to map research aims. The included articles were coded accordingly based on the objectives they aimed to address. Major themes, subthemes, and their corresponding codes and definitions were developed by the review team. A thematic analysis approach was similarly employed in the analysis of the gait assessment technologies.

## Results

### Selection of sources of evidence

Fig 1 shows the PRISMA flow diagram for the study’s selection method. A total of 729 articles were retrieved from three databases and other sources: 316 from Scopus, 280 from Web of Science Core Collection, 113 from CAB Abstracts, and 20 from the other sources. The de-duplication process left 474 records to screen. After screening abstracts and titles, we excluded 349 publications that were not related to field of bovine locomotion evaluation. By examining the remaining 125 in detail, a further 43 were excluded. Therefore, a final number of 82 articles were included in the scoping literature review.

### Characteristics of sources of evidence

The 82 included articles were published between 2000 and 2020, with the majority of articles (93%) published after 2005 (Fig 2). Datasets presented in the articles originated from 16 countries, with most originating from the United States (n = 16 articles; 19.5%), followed by Belgium and Canada (n = 12 articles; 14.6% and n = 11 articles; 13.4% respectively).

**Fig 2.**
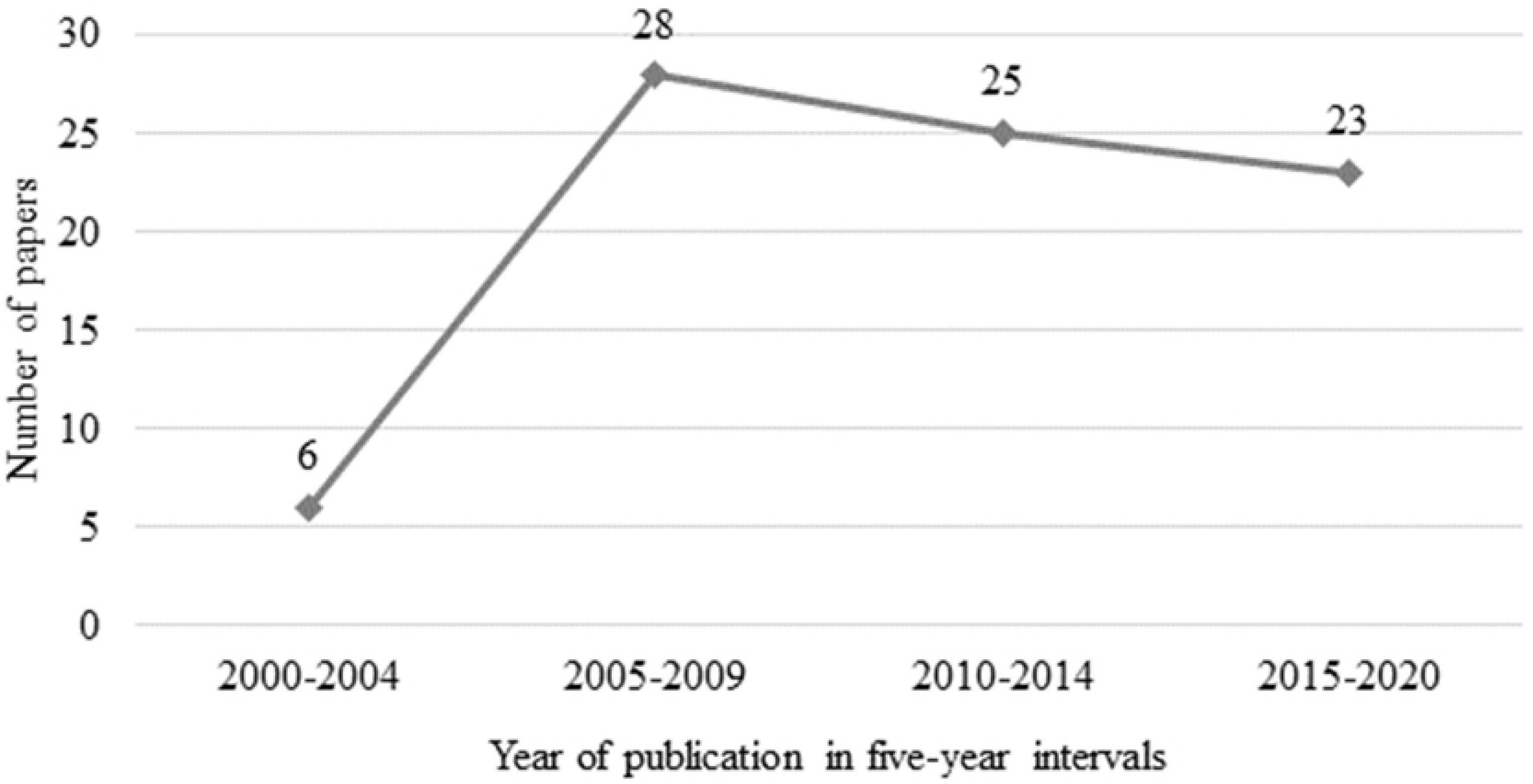
Number of peer-reviewed articles included in the scoping literature review over five-year intervals from 2000 to 2020.

Articles included in this review were published across 23 scientific journals. Journal of Dairy Science was the journal with the most articles included in this review (n = 33 articles; 40%), followed by Computers and Electronics in Agriculture (n = 10 articles; 12%).

### Thematic analysis of research aim

To understand the importance of using technology in bovine gait analysis, it is necessary to first examine the objectives of studies that have used these technologies. Therefore, a thematic analysis of research aims was conducted based on the stated objective(s) of the studies. Table 2 shows the thematic classification of research aims/themes, each theme was coded and described. There were four major themes: Gait/Claw biomechanics (GCB), Lameness detection (LD), Intervention/Comparison (IC), and System development (SD). Three of the major themes had subthemes. These were 1) Lameness detection: visual locomotion Scoring (LD-VLS), foot disorders (LD-FD), and automatic lameness detection systems (LD-ALDSs); 2) Intervention/Comparison: flooring type (IC-FT), hoof trimming (IC-HT), analgesics (IC-AN), and other (IC- Other); and 3) System development: development and validation (SD-DV), and system improvement (SD-IM).

**Table 2.**
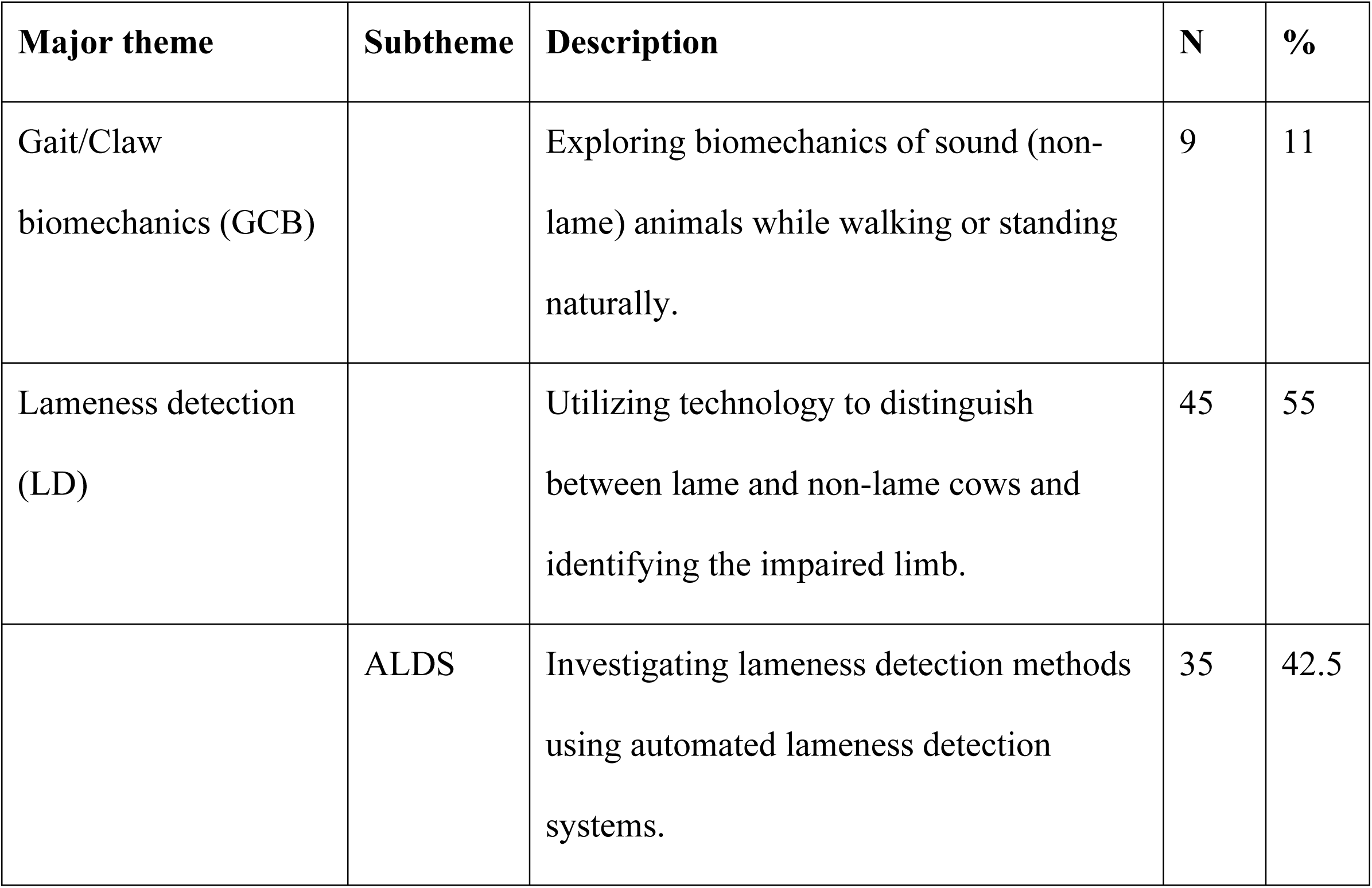
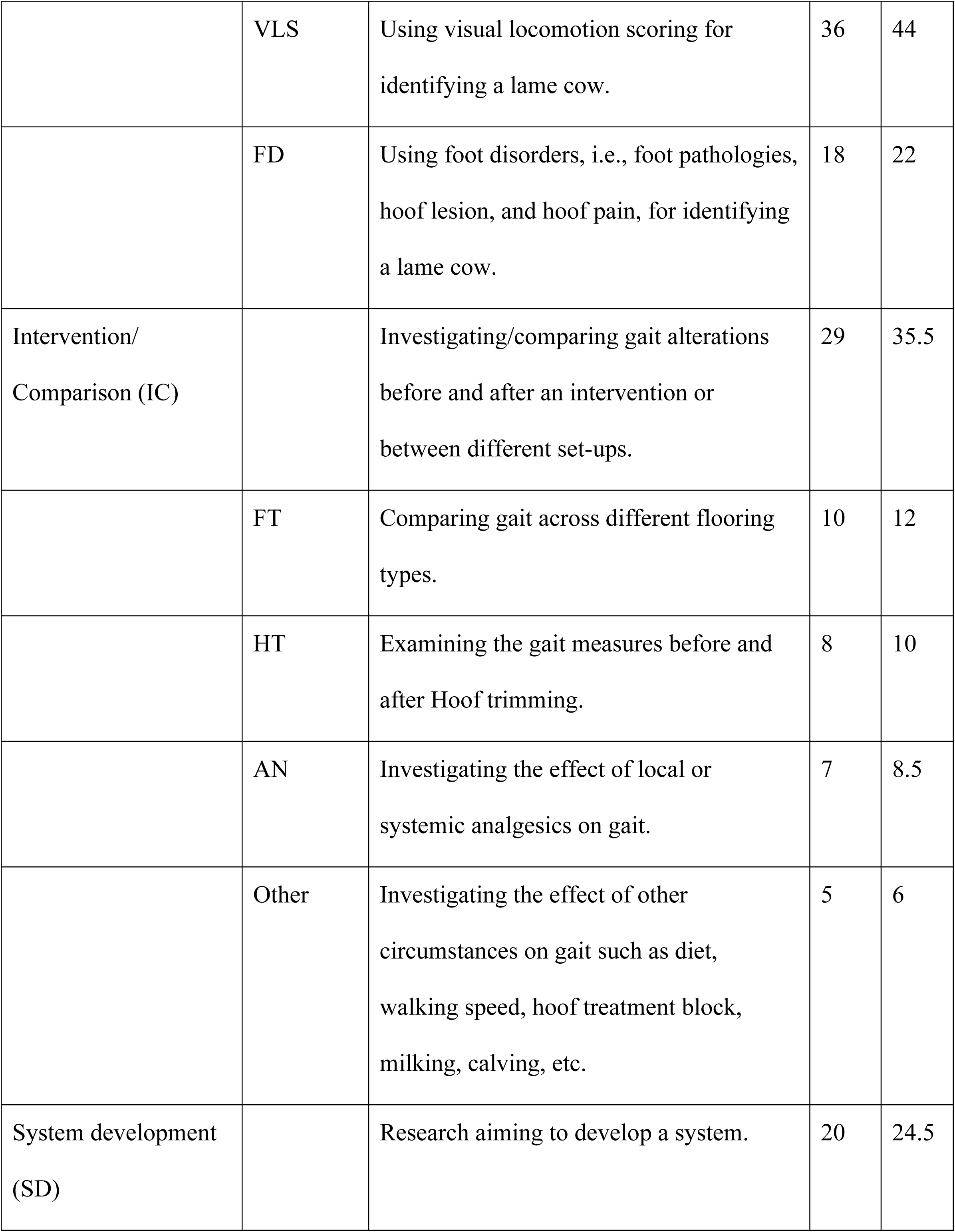
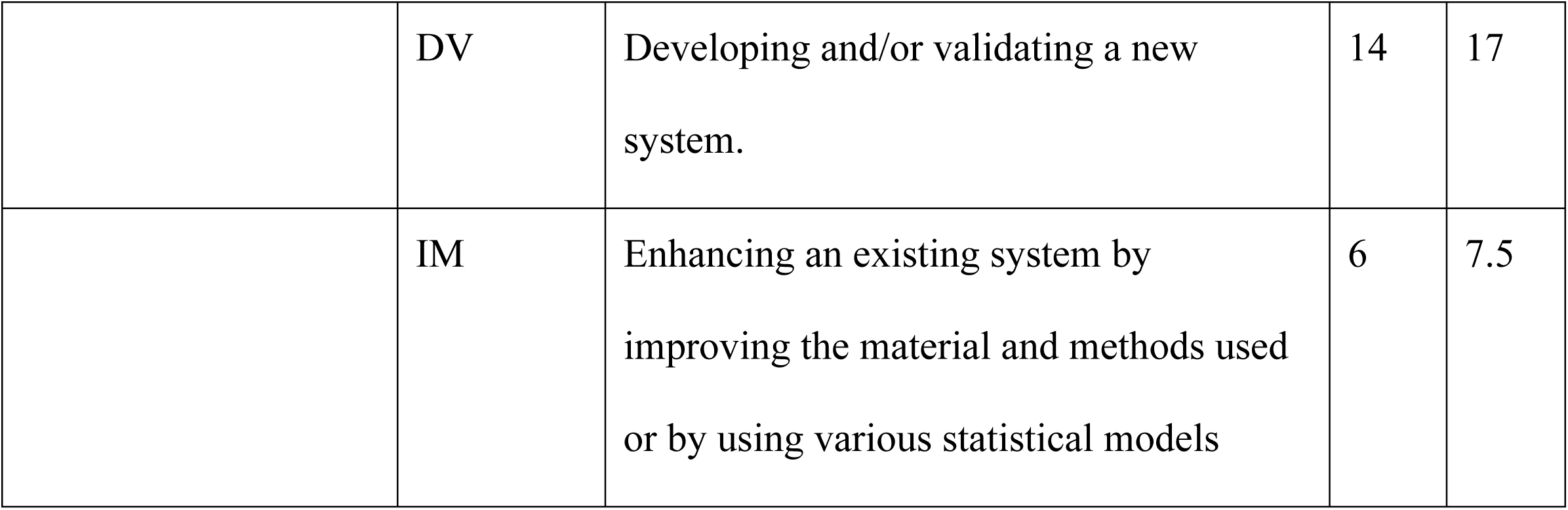
Published trends in bovine biomechanical research, with findings derived from thematic analysis of stated research aims of the 82 included articles. Some papers were categorized under more than one main theme or subtheme (i.e., they aimed more than one objective). Abbreviations: ALDS: automatic lameness detection system; VLS: visual locomotion scoring; FD: foot disorders; FT: flooring type; HT: hoof trimming; AN: analgesics; DV: development -validation; IM: improvement.

| Major theme | Subtheme | Description | N | % |
| --- | --- | --- | --- | --- |
| Gait/Claw biomechanics (GCB) |  | Exploring biomechanics of sound (non-lame) animals while walking or standing naturally. | 9 | 11 |
| Lameness detection (LD) |  | Utilizing technology to distinguish between lame and non-lame cows and identifying the impaired limb. | 45 | 55 |
|  | ALDS | Investigating lameness detection methods using automated lameness detection systems. | 35 | 42.5 |
|  | VLS | Using visual locomotion scoring for identifying a lame cow. | 36 | 44 |
|  | FD | Using foot disorders, i.e., foot pathologies, hoof lesion, and hoof pain, for identifying a lame cow. | 18 | 22 |
| Intervention/<br>Comparison (IC) |  | Investigating/comparing gait alterations before and after an intervention or between different set-ups. | 29 | 35.5 |
|  | FT | Comparing gait across different flooring types. | 10 | 12 |
|  | HT | Examining the gait measures before and after Hoof trimming. | 8 | 10 |
|  | AN | Investigating the effect of local or systemic analgesics on gait. | 7 | 8.5 |
|  | Other | Investigating the effect of other circumstances on gait such as diet, walking speed, hoof treatment block, milking, calving, etc. | 5 | 6 |
| System development<br>(SD) |  | Research aiming to develop a system. | 20 | 24.5 |
|  | DV | Developing and/or validating a new system. | 14 | 17 |
|  | IM | Enhancing an existing system by improving the material and methods used or by using various statistical models | 6 | 7.5 |

Lameness detection, i.e., identifying lame cows or an impaired limb using technology, was the most frequent research aim pursued in the gait analysis literature with 45 studies (55%, Table 2.4.1), followed by Intervention/Comparison studies (29 studies, 35.5%) and System development studies (20 studies, 24.5%). Studies that aimed to explore gait/claw biomechanics of a non-lame cow using a technology accounted for the lowest number of studies (9 studies, 11%) in the bovine gait analysis literature. The trend of four major research aims/themes (Fig 3) illustrates that lameness detection has maintained its dominance over the other research objectives for the last 15 years.

**Fig 3.**
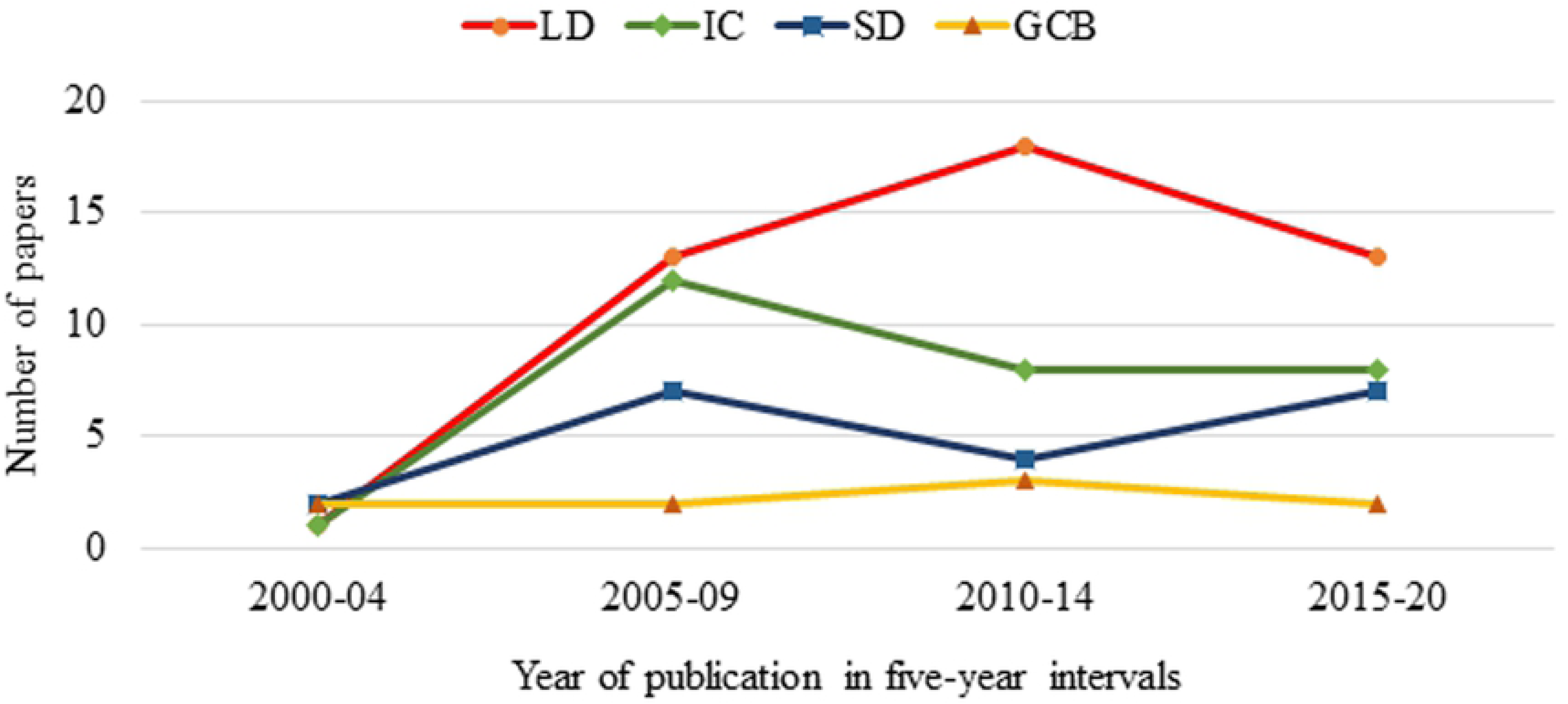
Research aims plotted over 5-year bins from 2000–2020. The four aims plotted were lameness detection (LD), intervention/comparison (IC), system development (SD), gait/claw biomechanics (GCB).

### Gait analysis technologies

Three main technologies employed in bovine gait analysis including force and pressure platforms (FPP), vision-based technologies (VB), and accelerometers. Table 3 shows an overview of these technologies and their subcategories that will be explained in detail further in this review.

**Table 3.**
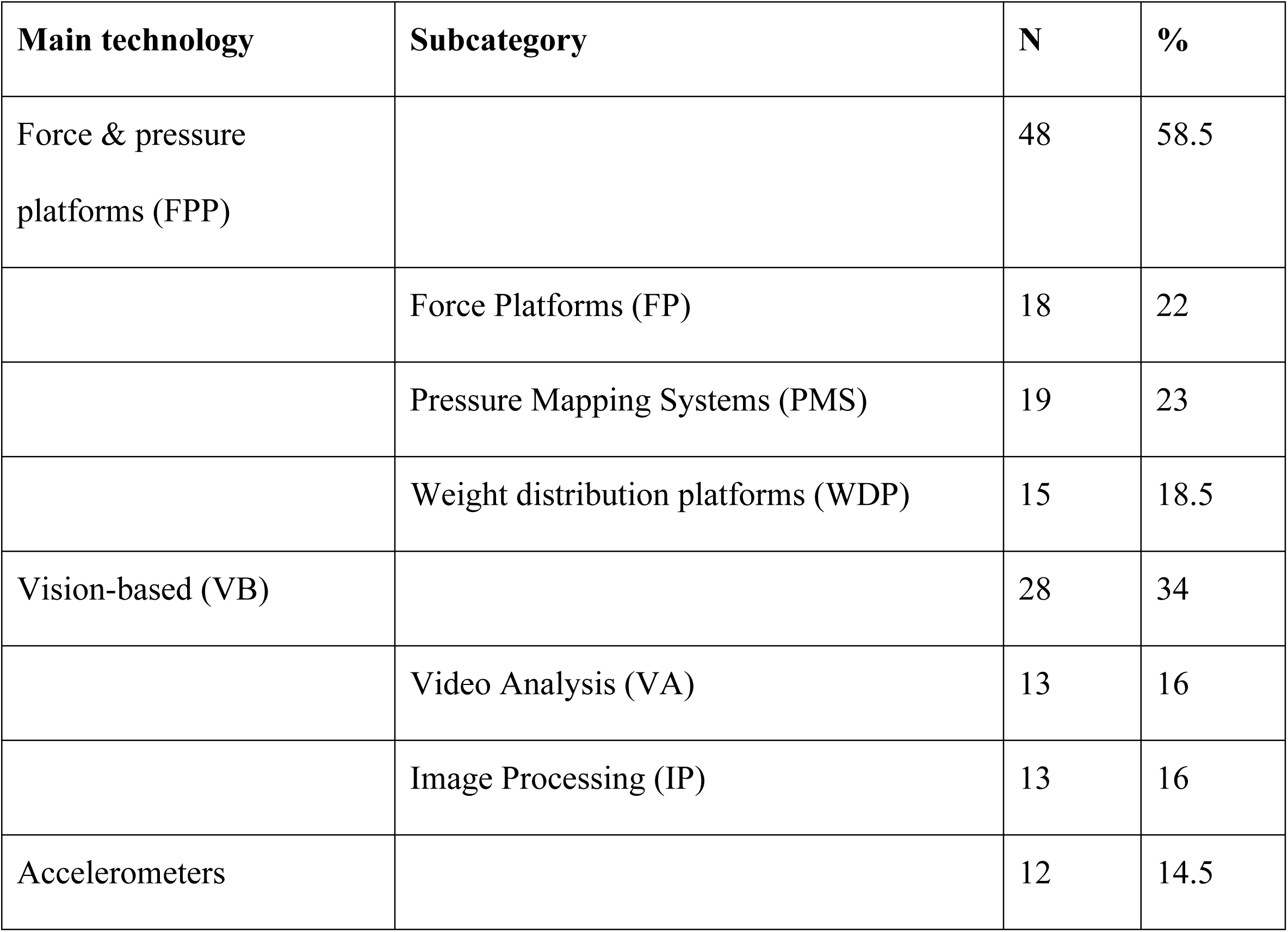
Three main gait analysis technologies and their subcategories.

#### Force and pressure platforms (FPP)

Force, pressure, and weight — along with their derivatives — are the main kinetic attributes measured by floor-sensitive plates. These measures are taken either when an animal is walking over the plates (dynamic measurements) or when standing on them (static measurements). Measuring plates were the most widely used technology (48 studies, 58.5%) in bovine biomechanical analysis. In the current review, these technologies are classified into three main groups: force platforms (FP), pressure mapping systems (PMS), and weight distribution platforms (WDP). More information on these three main kinetic technologies can be found in table 4.

**Table 4.**
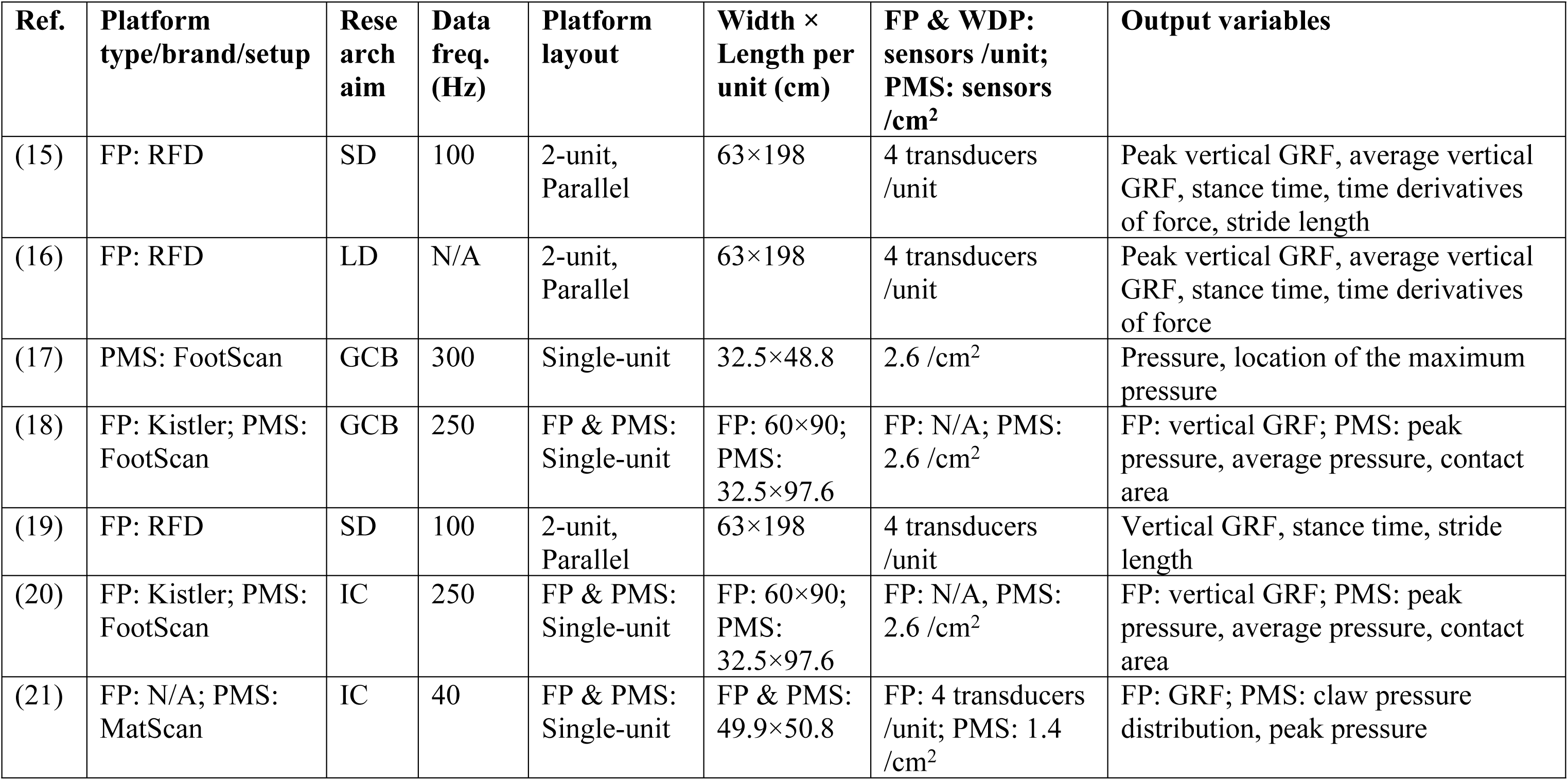
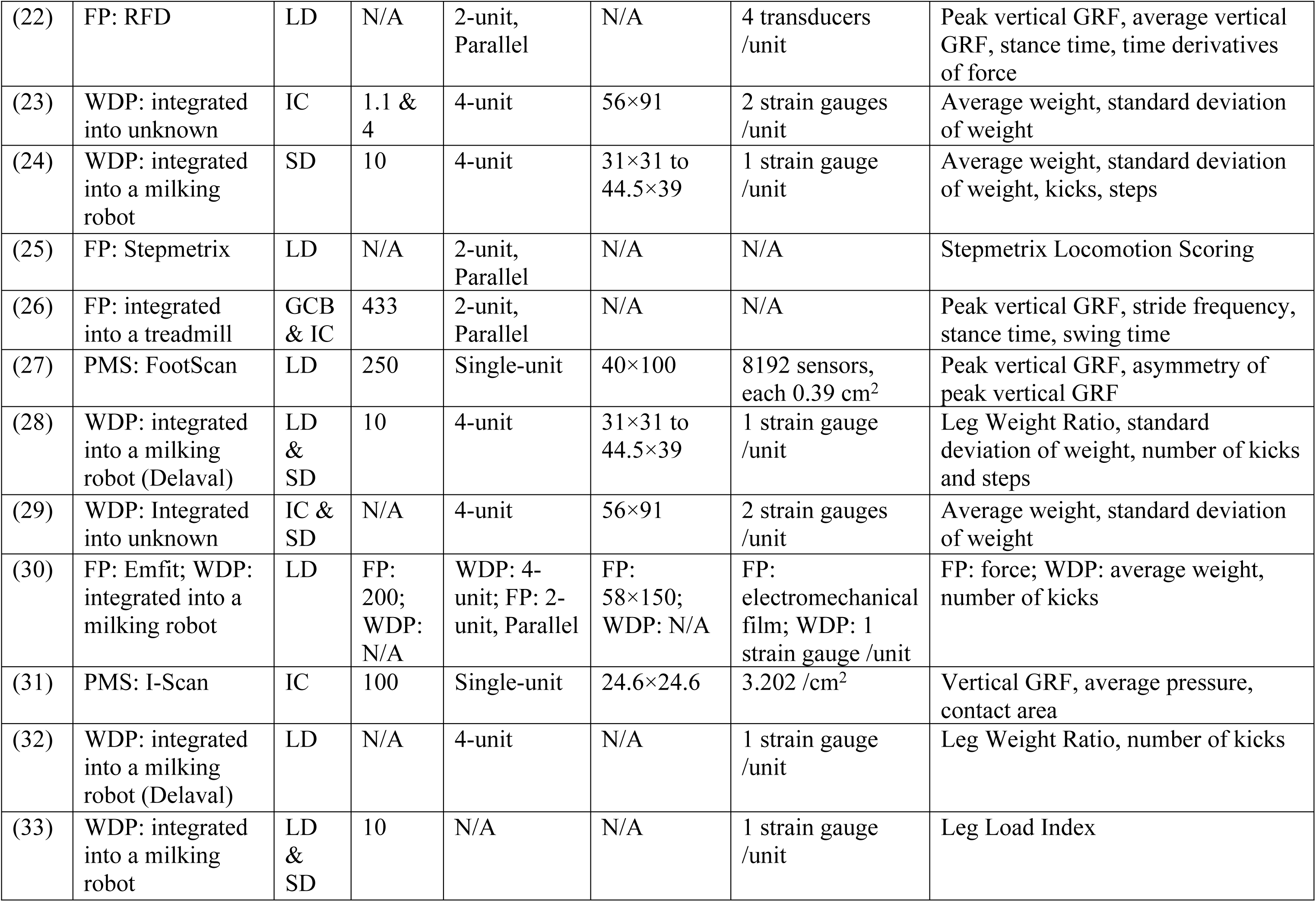
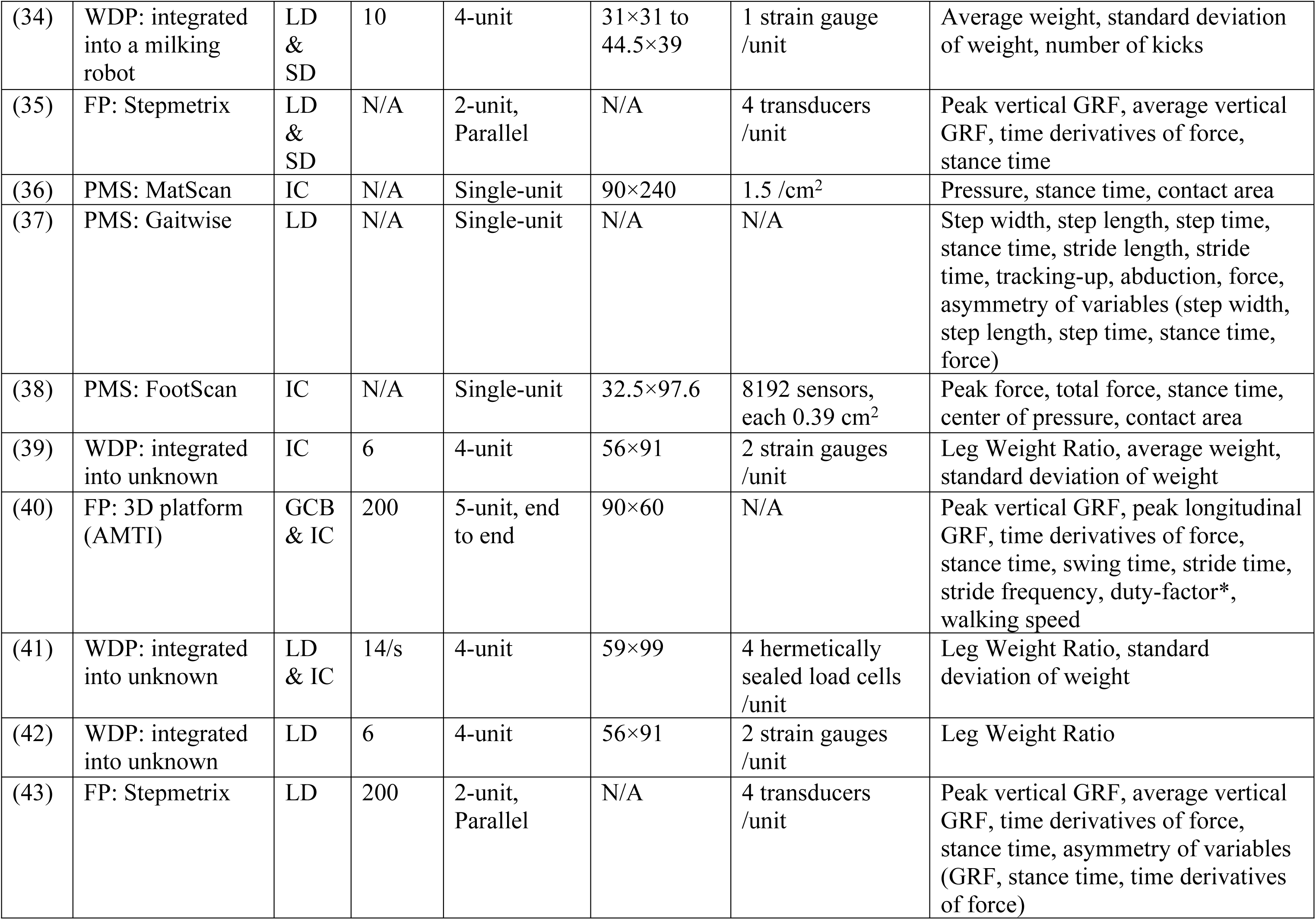
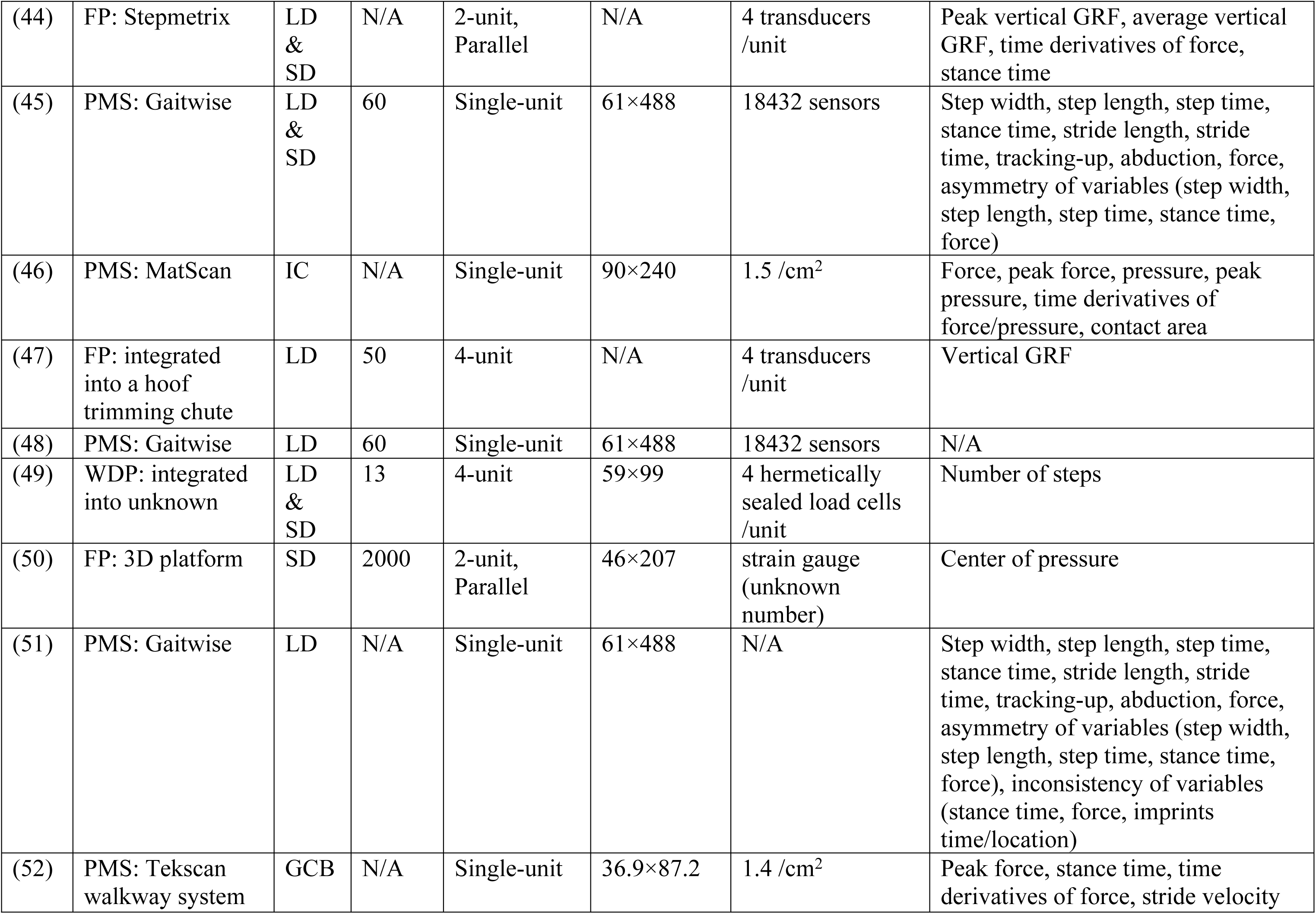
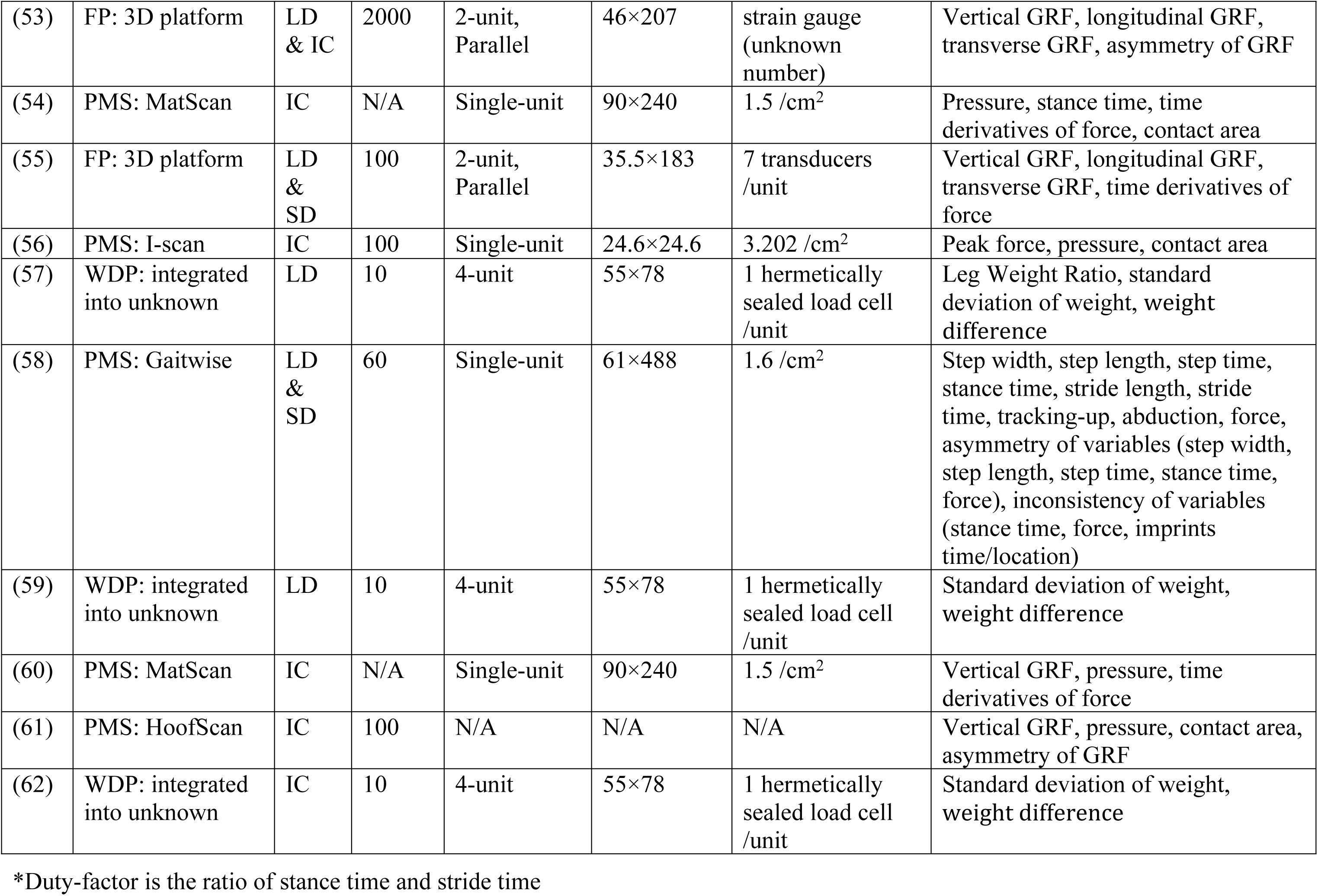
Overview of studies included in the review that used force and pressure platforms (FPP). Abbreviations: FP: force platforms; RFD: reaction force detection system; PMS: pressure mapping systems; N/A: not applicable; WDP: weight distribution platforms; SD: system development; LD: lameness detection; GCB: gait/claw biomechanics; IC: intervention/comparison; GRF: ground reaction force

##### Force platforms (FP)

Force platforms use one of several different types of transducers (aka load cells) which are installed under a cover plate to measure ground reaction forces (GRF) applied over the platform’s top surface. There were 18 bovine gait analysis studies that have used different force platform types, and these vary in the type and number of force transducers. The most common FP among the studies were 1D (single-axis) plates that can only measure the vertical forces. Alternatively, 4 studies included 3D force plates that can provide information on the three-dimensional components of force (vertical, transverse and longitudinal). One study also used a thin electromechanical film (Emfit) to measure the force generated as a result of a change in the film thickness (30). Additionally, a treadmill-integrated force measuring system equipped with 18 force transducers (9 on each side) was employed in one study to measure GRF while cows were walking at a controlled speed (26).

Most studies implemented a multiple-force-plate layout, usually a double setup with parallel orientation, to capture bi-lateral and/or individual footfall measurements of force. Three studies, however, used a single force plate system, which allowed for the evaluation of only one side of the animal (i.e., the fore and hind limbs of one side) during a measurement. The length of the FP varied between the studies that have reported platform dimensions, typically ranging from 0.9 to 2 m. An exception to this was the application of a 3-meter-long force platform system in a study that placed 5 smaller plates (each 0.6 m long) in a row to measure multiple consecutive footfalls (40). Force data recording rate also had considerable variation among the studies that reported it, ranging from 50 Hz to 2000 Hz. However, the usual recording frequency for most of the studies ranged from 100 to 250 Hz.

###### Force platform variables

Dynamic measurements recorded while animals walk over the plates are the primary way of using force platforms. However, measurements recorded while the animal is standing have also been used in two studies; one study used a force plate system comprised of four plates installed in a hoof-trimming chute (47), and the other used a combined system of force and pressure plates (20). Peak and average ground reaction forces (GRF) were the most important and frequent variables obtained from FP. The GRF measurements have been primarily measured vertically, except in the case of 3D platforms, which can supplement the vertical inputs with the transvers and the longitudinal inputs. In addition to GRF, other derivatives of force with respect to time (e.g., GRFω and impulse) have been measured. Centre of pressure (COP) and duty-factor (ratio of stance time and stride time) are other variables that have been measured using 3D force platforms. COP was measured to identify stance phases (50) or to detect left and right limbs (40). Stance time is the most frequent temporal variable that has been measured by most of the FP. More temporal variables such as swing time, and stride time were provided when a five-platform layout was used (40).

The studies that used Reaction Force Detection (RFD) systems (15) — Stepmetrix and the redesigned 3D version of the RFD system — usually presented their measurements as Limb Movement Variables (LMV), which are typically a collection of the following variables: peak GRF, average GRF, stance time, impulse, and GRFω. The Stepmetrix machine, as a lameness detection system, provides an automated locomotion scoring system (Stepmetrix locomotion scoring, SLS) using the LMVs. The SLS ranges from 1 to 100, and has been only reported in one study (25). Asymmetry measurements of some aforementioned variables such as GRF, stance time, impulse and GRFω have been also calculated by two studies (43, 53).

##### Pressure mapping systems (PMS)

Pressure mapping systems consist of a continuous network of sensors which measure multiple points of contact simultaneously, allowing for a detailed pressure map to be produced. They have been employed in 19 studies, 3 of which used a combined system of a pressure plate laid over a force plate. They typically come in the form of walkway plates and mats that allow animals to walk over or stand on them. However, one study developed an insole pressure system by attaching a sensor-equipped shoe (a leather claw shoe) to the cow’s left hind claw to measure kinetic outputs for both standing and walking conditions (61).

Pressure mapping systems had considerable variation in dimensions, ranging from 24.6 × 24.6 cm (I-Scan system to assess static pressure distribution of only the left hind foot) to 61 × 488 cm (Gaitwise system to capture more gait cycle steps for lameness detection purposes). The sensor resolution of a pressure mapping system is another aspect that varied among the studies that reported it, ranging from 1.4 to 3.2 sensing elements (pressure sensors) per cm^2^. Recording rate of data (frequency) also varied greatly among the studies, ranging from 40 to 300 frames per second (Hz).

###### Pressure mapping system variables

Kinetic outputs measured by PMS are typically obtained through dynamic assessments, with the exception of 4 studies that performed static measurements (17, 20, 31, 56). Vertical pressure, vertical GRF, impulse, contact area, and stance time are common variables measured among the studies using pressure mapping systems. One study also calculated center of pressure to investigate hoof pressure patterns during stance phase (38). Additionally, studies that have used the Gaitwise system (37, 45, 51, 58), provided mainly temporal (step time, stance time, and stride time) and spatial (step width, step length, stride length, tracking-up distance, and abduction) variables and their asymmetries. Relative force was the only force/pressure variable measured by the Gaitwise system.

##### Weight distribution platforms (WDP)

Four-scale weight distribution platforms consist of four independent recording units (one for each limb) that measure the weight distribution between limbs while the animal is standing. These platforms have been used in 15 studies, in which they were installed either inside or outside automatic milking systems. Studies incorporating WDP mainly aimed to detect leg problems, as lame cows reduce weight-bearing on the affected limb, consequently showing asymmetry in weight distribution across contralateral limbs (i.e., legs on the left vs. right side of the cow). Each recording unit typically contained 1 or 2 transducers (load cells) measuring the load applied on them. However, two studies used a weighing platform that had 4 load cells within each recording unit (41, 49). Dimensions of recording units ranged from 31 × 31 cm to 56 × 91 cm. The sizes of front and rear units may differ, as the rear recording units installed in the milking robots were usually larger than the front units. The duration of time that a cow stands on the weighing platform for data recording varied among studies, typically ranging from 2 to 5 minutes in 1 to 4 separate measurements. The reported recording rate (frequency) varied from 1.1 to 14 recordings per seconds (Hz).

###### Weigh distribution platform variables

Mean and standard deviation of the weight distributed between all legs or between a pair of legs were the most frequent variables that have been measured during the recording time in which animals stood on the weight distribution platform. Leg weight ratio (LWR) and weight difference (Δweight) were the other important variables associated with contralateral legs. The number and frequency of kicks and step behavior have been also calculated using weight distribution data.

#### Vision-based technologies (VB)

Vision-based technologies involve acquiring and analyzing the motion and posture (i.e., kinematic parameters) of a walking animal using videos or sequential images. These technologies have evolved considerably from manual annotation of images and videos to automatic motion/posture optical trackers, and computer vision and machine learning algorithms. The beginning of kinematic gait analysis in cattle using vision-based technology goes back to the study of Herlin and Drevemo (63), who measured angular patterns and hoof trajectories. Since 2000, there were 28 (34% of studies) published papers in bovine motion analysis that have utilized vision-based technology in different methods to measure locomotion variables while animals are walking through a corridor. Table 5 shows more information on vision-based technologies. The most frequent methods are video analysis (VA) and image processing (IP) techniques.

**Table 5.** Overview of studies included in the review that used vision-based technologies (VB). Abbreviations: N/A: not applicable; SD: system development; LD: lameness detection; GCB: gait/claw biomechanics; IC: intervention/comparison

| Ref. | Technique | Research Aim | Camera setup | Data freq. (Hz) | Number of recorded/analyzed strides/frames | Target anatomical regions | Output variables |
| --- | --- | --- | --- | --- | --- | --- | --- |
| (64) | Video analysis - Landmarks (markers) tracking | LD | 2D cam, side-view | 60 | 2 consecutive strides | Limbs | Stride length, max stride height, stride time, stance time, swing time, hoof speed, double and triple support, hoof trajectories |
| (65) | Video analysis - Landmarks (anatomical points) tracking | IC | 2D cam, side-view | N/A | 3 steps (2 strides) | Limbs & back posture | Walking speed, stride length, stepping rate, ROM (fetlock, Fetlock, Carpal and tarsal joints),backward angle horizontally (hoof, fetlock, Carpal and tarsal joints), back posture, Vertical fluctuation in the back |
| (66) | Video analysis - Landmarks (markers) tracking | IC | N/A | N/A | N/A | Limbs | Stride length, max stride height, stride duration, stance duration, swing duration, hoof speed, triple support |
| (67) | Video analysis - Landmarks (anatomical points) tracking | IC | 2D cam, side-view | 30 | 2 consecutive strides | Limbs | Stride length, ROM (Fetlock joints), stride time, stride velocity |
| (68) | Video analysis - Landmarks (markers) tracking | IC | 2D cam, side-view | N/A | 2 consecutive strides | Limbs | Stride length, max stride height, stride time, stance time, swing time, hoof speed, triple support, track-up |
| (26) | Video analysis - Hoof-ground contact order | GCB, IC | 2D cam (high-speed), side-view | 500 | 4 consecutive steps (3 strides) | Hooves | Ground contact order of claws and claw's zones |
| (69) | Video analysis - Landmarks (anatomical points) tracking | LD | 2D cam, side-view | 30 | 2 Strides | Hooves | Stride time |
| (70) | Image processing | LD | 2D cam, side-view | 15 | 2 Strides | Hooves | Track-up |
| (71) | Video analysis - Hoof-ground contact order | GCB | 2D cam (high-speed), side-view | 500 | 4 consecutive steps (3 strides) | Hooves | Ground contact order of claws and claw's zones |
| (72) | Image processing | LD | 2D cam, side-view | 15 & 20 | N/A | Hooves | Track-up |
| (73) | Image processing | LD | 2D cam, side-view | 30 | 4 frames (based on front/hind hoof-ground contact) | Back posture | Back posture curvature |
| (74) | Video analysis - Landmarks (markers) tracking | LD | 2D cam, side-view | N/A | N/A | Limbs | Stride length, track-up, ROM (fore fetlock, knee, hind fetlock, hock) |
| (75) | Video analysis - Landmarks (markers) tracking | GCB, IC | 2D cam, side-view | 15 | N/A | Limbs | Range of vertical and forward movement of limb joints |
| (48) | Image processing | LD | 2D cam, side-view | 20 | 1 step (right side) and 2 steps (left side) | Limbs | ROM of touch/release angles |
| (76) | Video analysis - Landmarks (markers) tracking | LD | 2D cam, side-view | N/A | N/A | Limbs & back posture | Stride length, track-up, Stride time, fetlock/hock/withers height, back length, hoof speed, head position |
| (77) | Image processing | LD | 2D cam, side-view | 25 | N/A | Back posture | Body movement pattern (BMP) |
| (78) | Image processing | LD | 3D cam (Kinect), overhead | 30 | Average 4.9 frames per video | Back posture | Back posture measure (BPM) |
| (79) | Image processing | LD | 2D cam, side-view & 3D Cam (Kinect), overhead | 25 (2D cam), 30 (3D cam) | Digital cam: 2 frames based on hind hoof-ground contact); 3D cam: average 2.56 frames per cow | Back posture | Back posture measure (BPM) |
| (80) | Video analysis - Landmarks (anatomical points) tracking | IC | N/A | N/A | 2 consecutive strides | Limbs | Walking speed, step length, stepping rate, stance time, track-up, support phase |
| (81) | Video analysis - Landmarks (markers) tracking | IC | 2D cam, side-view | 30 | 3-4 recorded strides, 2-3 analyzed strides | Limbs | Stride length, swing time, stance time, hoof height, number of steps, passage time, time per step. |
| (82) | Image processing | LD | 3D cam (Kinect), overhead | 30 | N/A | Back posture | Walking speed, back posture measure (BPM) and its variables |
| (83) | Image processing | LD | 3D cam (Asus), overhead | 30 | 2 consecutive strides ( average 70 frames) | Back posture (spine & hooks) | Height variation of hooks and spine regions |
| (84) | Image processing | LD | 3D cam (Kinect), overhead | 30 | N/A | Back posture | Body movement pattern (BMP) |
| (85) | Image processing | LD | 3D cam (Kinect), overhead | 30 | Average 5 frame per video | Back posture | Body movement pattern (BMP) |
| (86) | Image processing | LD | Integrated web cam, side-view | 25 | N/A | Limbs | Reduced speed, tracking up, asymmetry in gait and stance time, stride length, tenderness |
| (87) | Image processing | LD, SD | Dome cam, side-view | 25 | N/A | Limbs | Distribution of body pixels |
| (88) | High-speed fluoroscopic kinematography | GCB, IC | Biplane high-speed fluoroscopic | 500 | 1 stance phase | Limbs | ROM of interphalangeal joints |
| (89) | Object detection system | LD, SD | Dome cam, side-view | 25 | N/A | Limbs | Relative step size characteristic vector |

Video analysis (VA) techniques were used in 13 studies, eleven of which involved digitizing and tracking anatomical landmarks of a walking cow using motion analysis software to extract kinematic data either directly from a video (8 studies) or from consecutive still images obtained from a video (3 studies). Labelling and digitizing the anatomical landmarks can be done either manually, which requires manual localization of several points of interest, or using a marker-based system attaching reflective markers to palpable anatomical landmarks to measure kinematic data more accurately. Seven studies have employed a marker-based VA method using different types of passive markers (retroreflective markers) including reflective tape, color cardboard and reflective plastic balls. Additionally, two studies used the VA method to look at the pattern of ground contact of the claws of animals walking on a treadmill (26, 71). All VA studies employed only a single digital camera, which is usually positioned perpendicular to the plane of movement of the cows to record sagittal plane kinematics data. Therefore, they were only limited to analysis of two-dimensional (2D) kinematic data. Filming a walking cow has also been employed by a few studies to assist other technologies, i.e., force and pressure platforms and accelerometers, as a video synchronization in the determination of stride events (e.g., hoof strike, and toe off). These studies are not included under VB technologies in this review since they were not primarily used to produce gait variables.

Image processing (IP) is the other method of VB technologies that was used in 13 studies to analyze the motion and posture of a walking cow through an algorithm. After extracting still images of several desired frames from a video, some operations will be applied to the images to prepare them for the cow motion/posture recognition. Background subtraction is one of the major tasks in image processing procedure, which allows an image’s foreground, i.e., a moving cow, to be extracted from background for further processing. Then the motion/posture of the desired regions will be analyzed either manually or using machine learning algorithms. Studies that have employed IP techniques usually targeted one anatomical region. Back posture as one of the main lameness indicators is the most popular single target in the studies that utilized image processing techniques, followed by hoof and leg movement. Similar to the videography analysis method, the image processing studies also used a side-view camera, unless in the case of using 3D camera, i.e., depth-sensing camera, to produce depth images of the cow’s back posture that is positioned overhead.

The object/target detection system is a novel technique in the field of vision-based technology and computer vision that allows us to automatically identify and locate multiple objects/targets in a video. This technique, used recently in one bovine gait analysis study, works by applying deep learning algorithms to detect the leg, head, and back regions of the cow (89). However, only leg data has been analyzed and published. Biplane high-speed fluoroscopic kinematography (HFK) was another new technique to analyze bone movement with high accuracy in a 3D approach that has been utilized in a bovine study to evaluate the range of motion of interphalangeal joints (88).

Type of cameras varies among all the VB studies from a digital video camera to 3D depth sensors and dome cameras with various recording rates ranging from 15 to 60 fps. A high recording rate of 500 fps was set for the HFK study (88) as well as for two studies that walked the animals on a treadmill for video analysis (26, 71). In most of the studies it has also been stated that camera placement was set up in a position to allow researchers to evaluate at least two consecutive strides.

##### Vision-based technology variables

Vision-based technologies can provide distance (spatial) and time (temporal) parameters, angular range of motion (ROM) of joints, displacement, and velocities, all of which are classified under kinematic variables. Since video analysis techniques allow researchers to quantify motion and posture of multiple anatomical regions at the same time – i.e., several points (usually major joints) on fore and rear limbs, back posture, and head position – a more complete set of kinematic variables can be obtained. Stride length and stride time are the most frequent variables that have been measured using VA techniques, followed by maximum stride height, hoof velocity, stance time, swing time, triple support, hoof overlap and ROM of fetlock joint.

On the other hand, most of the studies that used image processing methods generally focused on one specific anatomical area leading to a particular variable. There are 7 IP studies that have focused only on the back posture and provided particular variables such as curvature angles (curvature angle of back around shoulders, curvature angle of back around hip joints, overall back curvature angle, curvature distance of shoulder and hip), back posture measure (BPM) and inverse radius. In addition to back curvature, one study (83) also looked at the hook bones in order to track the hind limb symmetry movements. There were two other studies that only looked at the hoof displacements and measured hoof location trackway, i.e., hoof overlap (70, 72).

Leg swing/movement has been another target for the kinematic studies using vision-based technologies. One study evaluated ROM of leg’s touch and release angles while another analyzed leg swing to generate six features referring to the gait asymmetry, speed, tracking up, stance time, and stride length (48, 86). Also, a different work adopted an object detection system to detect leg movement and measured the relative step size characteristic vector (89).

#### Accelerometers

Gait analysis in cattle has also been done using accelerometers which are attached to the cow’s leg to measure acceleration data. The output from accelerometers in bovine studies are generally classified as either behavior measures, including step activity, lying, and standing behaviors or as gait measures, including kinematics and kinetics of a gait cycle and symmetry between the legs. The current review focuses only on the gait measurement aspects of using accelerometers. There were 12 studies that have utilized the wearable sensors (i.e., 3D accelerometers) to measure gait variables. The details of these studies are shown in table 6.

**Table 6.** Overview of studies included in the review that used accelerometers. Abbreviations: Acc: Accelerometer; N/A: not applicable; WDP: weight distribution platforms; SD: system development; LD: lameness detection; GCB: gait/claw biomechanics; IC: intervention/comparison; IMU: inertial measurement unit

| Ref. | Acc type and combined tech. | Device info | Data freq. (Hz) | Research Aim | Target anatomical regions | Output variables |
| --- | --- | --- | --- | --- | --- | --- |
| (90) | 3D Acc | N/A | 25 | SD | 4 Acc on 4 limbs | Symmetry of variance acceleration |
| (91) | 3D Acc | Hobo Pendant | 33.3 | LD, IC | 4 Acc on 4 limbs (fetlock) & 1 Acc on thoracic belt | Acceleration variables, walking speed |
| (75) | 3D Acc & Cinematography | N/A | 200 | GCB, IC | 4 Acc on 4 limbs | Vertical, lateral and forward movement |
| (92) | 3D Acc | RumiWatch | 10 | SD | 1 Acc on one of the hind limbs | Stride time, stride length |
| (93) | 3D Acc | RumiWatch | N/A | LD | 2 Acc on hind limbs | Stride time, stride length, walking speed |
| (81) | 3D Acc & Cinematography | Hobo Pendant | 33 | IC | 2 Acc on right limbs (front & hind fetlocks) | Mean acceleration, asymmetry of variance acceleration |
| (94) | 3D Acc, IMU* & Cinematography | N/A | 400 | LD, GCB | 1 Acc on one if the hind limbs & 2 Acc on both hind limbs | kinematic outcome (gait cycle, stance time, swing time), and kinetic outcome (foot load, toe-off) |
| (95) | 3D Acc & Cinematography | N/A | 400 | SD | 2 Acc on both hind limbs | kinematic outcome (gait cycle, stance time, swing time), and kinetic outcome (foot load, toe-off) |
| (96) | 3D Acc & Cinematography | N/A | 400 | IC | 2 Acc on both hind limbs | kinematic outcome (gait cycle, stance time, swing time), and kinetic outcome (foot load, toe-off) |
| (59) | 3D Acc & WDP | N/A | 400 | LD | 2 Acc on affected (lame) front/hind limb pair | kinematic outcome (gait cycle, stance time, swing time), and kinetic outcome (foot load, toe-off) |
| (97) | 3D Acc | N/A | 100 | IC, SD | 1 Acc on one of the hind limbs | Stride segmentation |
| (62) | 3D Acc & WDP | N/A | 400 | IC | 2 Acc on affected (lame) front/hind limb pair | kinematic outcome (gait cycle, stance time, swing time), and kinetic outcome (foot load, toe-off) |

\*IMU is a device that integrates 3D accelerometers, 3D gyroscopes, and a magnetometer

The typical location for attaching the accelerometer is above the fetlock joint at the metatarsal/metacarpal level of a limb; however, 2 studies placed accelerometer at thoracic vertebrae level as well. The main objective of using an accelerometer is to detect a lame cow or to find a gait abnormality in intervention studies. The number of accelerometers per cows depends on the research set-up and the purposes. Studies that used only one accelerometer per cow, i.e., attached to one of the hind limbs or the affected (lame) limb, looked at the consistency/inconsistency of a variable within an animal or the differences between animals. In cases where more than one accelerometer per animal was used, accelerometers were worn on 2 limbs (hind limbs only or one side of the cow) or all 4 limbs to study differences in acceleration data between the limbs. The sampling rate of the accelerometer varies among the studies, ranging from 10 to 400 Hz. It stands to reason that the studies which employed accelerometers with high sampling rates (more than 100 Hz) could provide more details on events and moments of a gait cycle.

##### Accelerometer variables

Measuring asymmetries and differences of acceleration data across the legs is one of the main parameters in the studies which employed more than one accelerometer per cow. Acceleration data has also been used to extrapolate kinematic and kinetic variables. Studies that used low frequency (less than 40 Hz) accelerometers have estimated stride time, stride length and walking speed in addition to behavioral measurements, while accelerometers with higher sampling rates have measured more detailed parameters of a gait cycle including stance and swing duration as temporal kinematic outcomes and foot-load, heel-off, and toe-off as kinetic outcomes.

### Gait technology trends

The trend of technology use in bovine gait analysis has changed throughout the last two decades (Fig 4). The use of both FP and PMS was the starting point of gait analysis research in bovine between 2000 and 2004, while the use of IP techniques of vision-based technologies and accelerometers have grown considerably over the last decade and have surpassed the other technologies.

**Fig 4.**
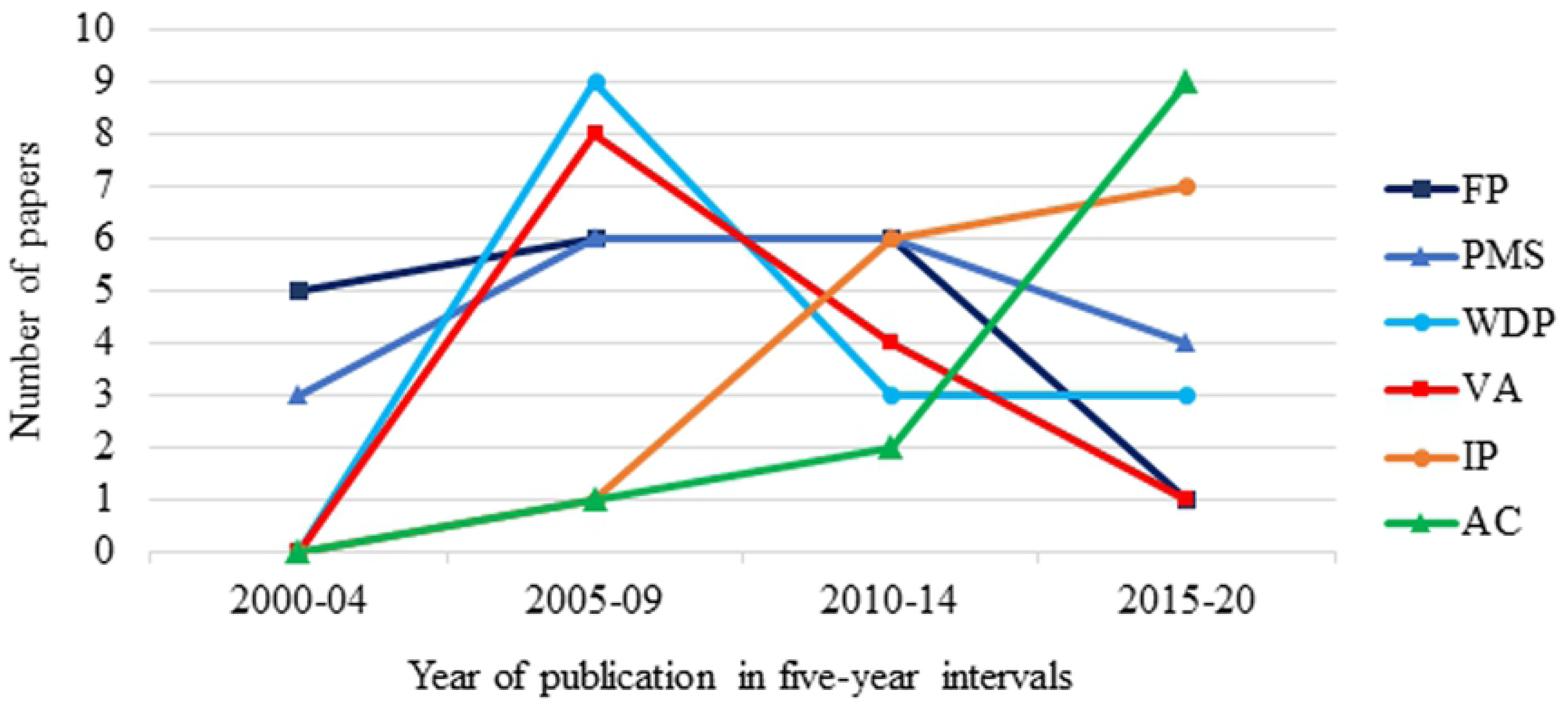
Technology trends in bovine gait analysis plotted in 5-year bins from 2000–2020. Force platforms (FP), pressure mapping system (PMS), weight distribution platforms (WDP), video analysis (VA), image processing (IP), and accelerometer (AC).

## Discussion

The field of bovine gait analysis stems from equine gait analysis and human movement research, adapting technologies that initially were developed in horses or humans such as force and pressure platforms, video analysis techniques, and wearable sensors. Theoretically, the research on equine locomotion can be easily applied to cows, since both species are considered large quadrupeds with many similarities in locomotion traits. Our review was able to include a total of 82 published papers related to objective gait analysis in bovine studies, compared to 510 papers (432 papers with the same period, between 2000 and 2018) included in a review with the same scope in equine gait analysis literature (98). This confirms that gait analysis in bovine research and practice is less covered compared to equine. The current main roles of the horse as a sport and leisure animal and the importance of its locomotor system performance is possibly the reason behind the difference in representations in scientific literature (99). Moreover, equine quantitative gait analysis is not only limited to lameness studies; with many other practical uses in clinics and training centers with the goal of improving equine performance, optimizing training, and enhancing the horse-rider relationship (98, 100). In cows, however, as a livestock animal, lameness and identifying impaired locomotion remains the primary topics investigated using gait analysis technologies. Indeed, lameness is an important issue in dairy cattle and is one of top 3 reasons for involuntary culling cows in North-America (101), thus the use of automated technologies for early detection and prevention is a main focus of interest. The results of this review also confirm that lameness detection is the main purpose of gait analysis research on cows (55% of studies) and the efforts for automatizing lameness detection on farms have influenced the research trend. Even most of the intervention/comparison (IC) studies looked at identifying impaired limb and gait abnormalities between different set-ups, confirming the dominance of lameness detection in gait analysis. Nevertheless, there is a limited number of studies on exploring natural (i.e., when cows are sound) gait/claw biomechanics using various technologies (11% of studies), especially between different breeds and size of cows, and investigating gait at early life. In fact, with regard to early life gait research, there is only one study that looked at the gait kinetics of 4-6 months old male calves. These are areas that could be the subject of future studies. While over a third of the literature is on IC studies (35.5%), the impact of other factors of welfare concerns and cow management (e.g., housing systems, stall dimensions, heat stress, outdoor access) on cow’s locomotion using gait analysis technology are seldom studied. To the best of our knowledge, there is no study published on the use of gait analysis technologies in beef cattle despite the fact that lameness is also a concerning issue in beef industry (102).

### Strengths and weaknesses of cow gait technologies

#### Force Platforms

Force platforms (FP) are considered the gold standard because of their recognized accuracy and high frequency measurements. Beyond vertical force, 3D force platforms can output shear forces (i.e., transverse, and longitudinal forces). This allows researchers to investigate more in depth the gait attributes affecting transverse and longitudinal components of force (55). Nonetheless, one of the FP limitations is that it is challenging to measure individual footfalls when more than one foot touches the plate at the same time (i.e., the best results are achieved when only one foot fits on the plate). Based on this, force plates generally come in multiple-platform layouts to register forces exerted by each hoof strike. However, walking animals on a multiple-platform setting to fit footfalls on the designated plates is challenging per se.

Research aim analysis shows that efforts have been made to employ FP as an automatic lameness detection system, generally using a commercial version of force platform for on-farm use (i.e., StepMetrix™, Boumatic LLC, WI, USA). Despite this, the use of FP in studies dropped since 2015, suggesting that the weaknesses of FP may outweigh the strengths, especially as other technologies, by comparison to FP, are capable of measuring a more diverse assortment of variables that can more robustly analyze gait.

#### Pressure Mapping Systems

Unlike the FP, pressure mapping systems (PMS) can distinguish and segment individual footprints regardless of how many feet are on the ground. As such, they can be used as a single mat or plate in different dimensions which makes them more applicable for kinetic analysis in quadrupeds. PMS can also quantify a wide range of pressure/force (only in vertical dimension), temporal, and spatial parameters, depending on the device’s specifications. Research aim analysis shows that PMS have been utilized to address different aims including evaluation the effects of hoof trimming, local analgesics, and flooring type on cows/claws’ biomechanics. In addition, lameness detection is another research aim that was studied, particularly using a longer version of pressure platforms (i.e., Gaitwise system). Utilizing a longer pressure mapping system allows registering more gait cycles and steps during a passage, which is a key for lameness detection purposes. In comparison with FP, PMSs are more practical and efficient in terms of easy implementation and generating more diverse outcomes either in research studies or at the farm level.

#### Weight Distribution Platforms

While being a limitation for FP and PMS systems as well, numerous failed measurements because of placing hooves outside of the measuring zones have been reported as the primary weakness for WDP. WDP measures variables only while an animal is standing, so the outcomes are limited to static measures and no information on gait cycle kinetics can be taken. Nevertheless, the strength of WDP is the fact that it is not just a quick snapshot of the cow walking, but a longer review of how she distributes her weight. A lack of even distribution of weight makes it easier to pinpoint the limb of issue in these cows with greater certainty, as the reluctance to bear weight in this case is usually a sustained measure. Moreover, they can be easily implemented in farms with the least space occupation, such as embedded in the flooring of milking robots or cattle chute. The results of our scoping review confirm this, since WDPs are primarily used for lameness detection purposes at the farm level.

#### Video Analysis

Video analysis is the gold standard for kinematic measurements. They are able to provide a variety of gait variables from distances to ROM, velocities, and trajectories. They allow researchers to investigate multiple anatomical locations on the body with a single passage, yielding a better understanding of different aspects of an animal locomotion. Since impaired locomotion (lameness) can manifest in different ways (e.g., from short stride length to back arch, joint stiffness, asymmetries), a system that can consider all of these variables can provide a more accurate analysis. However, these systems, especially marker-based VA systems, pose several limitations, including marker placement error, observer (digitizer) error, and the time-consuming process of manual/semi-manual tracking. Moreover, it requires animal preparation (attaching markers on animal’s body) and a specific environment (room) wherein lighting conditions can be controlled and ample space can be provided so as to not block the cameras’ views of the markers. All these procedures can be challenging and make these techniques unfeasible to implement on commercial farms. The results of this review illustrated that the use of VA techniques have decreased over time, from 8 studies between 2005 and 2009 to only 1 study over the last five years.

Moreover, all the previous studies that used VA technology were limited only to 2D kinematic analysis which includes several limitations by comparison to three-dimensional (3D) analysis, where multiple synchronized cameras are being used. Some of these limitations include parallax error and perspective error that occur when subjects move away from the optical axis of the camera and subjects are restricted to movement within the plane of calibration and accuracy is compromised. Also, 2D angular data are limited to one degree of freedom (DOF) for joint rotation within the sagittal plane, while 3D kinematic data allow the researcher to explore the full range of motion (ROM) and orientation of segments and joints within 3D space (103). Therefore, although the use of VA techniques is challenging, the high accuracy of these systems, especially the 3D video analysis systems, makes them a reliable reference for the future works of using marker-less VA technologies.

#### Image Processing

By comparison to VA systems, the downside of the IP methods is that, since they usually focus on one anatomical region as a lameness indicator, their outcomes are limited to the targeted region and the other facets of lameness will be missed. However, use of machine learning algorithms and eliminating the requirement of animal preparation (e.g., with reflective markers) allows them to be utilized as automated systems on commercial farms. Also, the use of overhead 3D depth image cameras instead of the side-view digital cameras benefited this technology to overcome the challenges of side-view image capturing at farm conditions, including occlusions and sensitivity to lighting variance in order to facilitate the image segmentation of the foreground from the background. Our results showed that efforts for implementing IP technology as automated lameness detection systems on farms has become more prevalent in gait analysis studies over the last decade (13 studies between 2010 and 2020).

#### Accelerometers

Accelerometers have been used as a technology for gait analysis in cattle studies since 2009 and have been the most widely used technology in the last 5 years. Previous studies showed that pedometers (accelerometers) can be a promising tool for detecting lameness and foot pathologies in dairy cows. Uncontrolled walking speed and cow traffic as well as behavior are the usual limitations reported for gait analysis using FP, PMS and WDP and vision-based technologies, but not for accelerometers. Accelerometer outcomes are not dependent on a single passage or a specific walking corridor. At present, however, accelerometers still carry limitations. For example, not all variables can be measured using a single pedometer, with useful gait metrics for detecting impaired gait, such as asymmetry of acceleration between legs, requiring 2 or more devices. Technologies that require more hardware can be a barrier to adoption on commercial farms (104). Moreover, current pedometers used on farm generally have lower data sampling rates as they are less costly than high-frequency pedometers; however, this trade-off can compromise the efficacy of the pedometer use for detecting impaired gait. Low frequency accelerometers can make gait analysis more challenging and requires more research.

### Limitations

There are a few limitations to this review. Only studies using technology for direct kinetic and kinematic gait analysis were included. This excludes 1) studies that used technology to evaluate gait based on cow’s behavior and 2) studies that did not measure kinetic and kinematic parameters (e.g., a study utilized cows footfall sound volume to detect claw lesions (105)). In addition, including only in vivo studies that conducted research only on live animals, has left out the studies that measured gait variables using cow’s foot specimen. Furthermore, the results of this review did not reflect information regarding study limitations because most of the studies have not stated their limitations.

## Conclusion

A strong demand for automatic lameness detection has influenced the path of development for quantitative gait analysis technologies in bovine. Although the field of bovine gait analysis has evolved considerably through the last two decades, more research is needed to achieve more accurate, practical, and user-friendly technologies for research and on-farm applications. Force and pressure platforms, image processing techniques, and accelerometers have shown that they are capable and practical methods for automated lameness detection systems, despite their weaknesses. Nevertheless, computer vision technologies using deep learning and wearable sensors (e.g., accelerometers) are promising emergent technologies that could be useful for the next stage of quantitative gait analysis in bovine.

